# Early development of electrophysiological activity: contribution of periodic and aperiodic components of the EEG signal

**DOI:** 10.1101/2022.10.04.510765

**Authors:** Josué Rico-Picó, Sebastián Moyano, Ángela Conejero, Ángela Hoyo, M. Ángeles Ballesteros-Duperón, M. Rosario Rueda

**Author notes:** Corresponding author MSc. Josué Rico-Picó, Adress: Cuesta del Hospicio, S/N, Facultad de Psicología, 18071 (Granada, Spain).

## Abstract

Brain function rapidly changes in the first two years of life. In the last decades, resting-state EEG has been widely used to explore those changes. Previous studies have focused on the relative power of the signal in canonical frequency bands (i.e., theta, alpha, beta, gamma). However, EEG power is a mixture of a 1/f-like background power (aperiodic) in combination with narrow peaks that appear over that curve (periodic activity; e.g., alpha peak). Therefore, the relative power may capture both, aperiodic and periodic brain activity, misleading actual periodic changes in infancy. For this reason, we explored the early developmental trajectory of the relative power in the canonical frequency bands from infancy to toddlerhood and compared it to changes in periodic activity in a longitudinal study with three waves of data collection at ages 6, 9, and 16 to 18 months. Finally, we analyzed whether the periodic activity and/or aperiodic components of the EEG contributed to explaining age changes in relative power. We found that relative power and periodic activity trajectories differed in this period in all the frequency bands but alpha, and we replicated an increment of alpha peak frequency. We found that age changes in aperiodic parameters (exponent and offset) depend on the frequency range. More importantly, only alpha relative power was directly related to periodic activity but other frequency bands were predicted also by aperiodic components. This suggests that relative power is capturing the developmental changes of the aperiodic brain activity and, therefore, more fine-grained measures are needed.

---

Great changes in the structure and functional brain activity take place in the first years of life, which has been related to the emergence and development of diverse cognitive processes (Gabard-Durnam & McLaughlin, 2020; Gilmore et al., 2018). Among all the techniques that measure brain function, electroencephalography (EEG) has been widely used to characterize functional development in infants and toddlers due to the easiness and adaptability of its application (Saby & Marshall, 2012).

The EEG signal offers information about neural oscillations, reflecting the synchronization and desynchronization of neuronal activation at different rhythms in both micro and macro neural circuits (Buzsáki & Draguhn, 2004; Buzsáki, 2006; Cohen, 2017; see Buzsáki, Anastassiou & Koch, 2012 for a review). Baseline EEG activity is usually registered at rest (resting-state EEG; rs-EEG), and it offers information about the intrinsic activity of the brain without the constraints of a task. As a result, the rs-EEG has been widely implemented to explore brain function development, as it does not require following particular task instructions. In infants, rs-EEG protocols usually consist of paying attention to an external stimulus (e.g., soap bubbles) to make the infants as calm as possible (Anderson & Perone, 2018; Bell, 1998; Saby & Marshall, 2012).

Resting state EEG provides several measures, ranging from signal energy to connectivity measures. However, the gold-standard measurement in infant rs-EEG consists of decomposing the signal to extract the power in canonical frequency bands: theta (3 – 6 Hz), alpha (6 – 9 Hz), beta (9 – 20 Hz), and gamma (20 – 45 Hz; Saby & Marshall, 2012). The power of each frequency band can be obtained in absolute terms when keeping the original values, or relative when the energy of a particular frequency is divided by the power of the rest of the signal or related to other frequency bands (e.g., theta-beta ratio; Anderson & Perone, 2018; Saby & Marshall, 2012; Trujillo et al., 2019).

When rs-EEG has been explored along with the development of the first years of life, the relative power is more sensitive than absolute power because absolute power can be affected by skull changes over the lifespan (e.g., Marshall et al., 2002). Indeed, the transition from infancy to toddlerhood seems to be a period of rapid reconfiguration of relative power in all the frequency bands. Firstly, there is evidence that theta relative power diminishes in infants compared to older children. Secondly, the alpha peak emerges around the fourth month of life. It appears as a sudden energy “bump” between 5 and 7 Hz that shifts towards higher frequencies and augments its relative power in the first years of life (Gasser et al., 1988; MacNeill et al., 2018; Orekhova et al., 2006; Stroganova et al., 1999; Whedon et al., 2020). Furthermore, alpha band relative power correlates between early and follow-up sessions in infants, which accounts for the intra-individual stability of this measure (Marshall et al., 2002). Finally, although research on higher frequency bands in infants is still scarce, a study by Tierney and colleagues (2012) suggests a reduction of frontal beta and gamma between the fifth month and the second year of life. In addition, the relative power in different frequency bands is related to individual differences in cognitive development (Anderson & Perone, 2018; Bell & Wolfe, 2007; Benasich et al., 2008; MacNeill et al., 2018; Whedon et al., 2020) and in infants at risk of neurodevelopmental disorder (Arns et al., 2013; Begum-Ali et al., 2022; Gabard-Durnam et al., 2019; Tierney et al., 2012), which signal its relevance.

Although previous research suggests that relative power is sensitive to developmental changes, it only considers the energy of canonical frequency bands that do not separate the background activity (aperiodic) from the periodic brain activity (Donoghue, Haller, et al., 2020; He, 2014; Ostlund et al., 2022; Voytek et al., 2015). In fact, the EEG power is a composite of a 1/f-like curve that accounts for most of the signal (aperiodic) with some “bumps” or peaks that appear over it (periodic or oscillatory). More specifically, the aperiodic background curve has been defined with two parameters: offset (power at the lowest frequency of the aperiodic curve) and the slope or exponent of the curve. On the other hand, periodic brain activity has been focused on the frequency of the peak and the energy above that curve.

As a result of this conceptualization of EEG power, some authors argue that developmental variations of absolute and relative power can be due to changes in either aperiodic or periodic components of the EEG (Donoghue, Dominguez, et al., 2020; Ostlund et al., 2022; Samaha & Cohen, 2022) Consequently, different research groups have tested the maturation of aperiodic and periodic components maturation (e.g., Cillier et al., 2021). Regarding infancy, recent research has shown a reduction in the slope from the zeroth to the seventh month of life (Schaworonkow & Voytek, 2021). This pattern seems to be constant as the background curve flattens from age 3years-old onward (Donoghue, et al., 2020b; Hill et al., 2022; McSweeney et al., 2021; Voytek et al., 2015; Voytek & Knight, 2015). Interestingly, the aperiodic parameters have been linked to cognitive processes and neurodevelopmental disorders, which speak for their biological relevance (Demuru & Fraschini, 2020; Immink et al., 2021; Karalunas et al., 2022; Shuffrey et al., 2022). On the other hand, developmental studies about periodic activity have mainly focused on the alpha frequency band. In infants, Schaworonkow & Voytek (2021) found that alpha frequency and the number of alpha burst increased in the first years of life, which also occur during childhood (e.g., Cellier et al., 2021).

The previous research points out that relative power development maturation is parallel to changes in periodic and aperiodic components of the EEG. Consequently, aperiodic and periodic components might underly the changes in relative power. Thus, further research is needed to unveil the relationship between relative power and the aperiodic and periodic components, as well as the differential trajectories and stability of the signal in unexplored periods of the lifespan, such as the transition between infancy to toddlerhood. For this reason, we registered rs-EEG in a longitudinal cohort of babies who were examined in three different waves (6, 9, and 16 to 18 months of age) and computed the relative power of canonical frequency bands, periodic brain activity, as well as the offset and exponent of the background curve. Our objective was to compare the trajectories of relative power and periodic brain activity and age-related changes in the aperiodic components. Also, we tested the intra-subject stability of the signal and analyzed whether the relative power was related to the aperiodic components and periodic power. We expected to replicate changes in relative power previously found in studies during infancy. That is, a reduction of relative power in theta, beta, and gamma, but an increase of alpha’s relative power. In addition, we expected similar trajectories of the periodic brain activity and a reduction of the exponent and offset aperiodic components with age. Also, we predicted that the measurements across waves would be correlated, which would mean that rs-EEG measurements are stable in the transition between infancy and toddlerhood. Finally, we hypothesized that the relative power would be related to both periodic power and aperiodic components. In such a case, it would suggest that relative power conflates background and actual periodic brain activity and would be relevant to interpret future rs-EEG studies.

## 2. METHOD

### 2.1. Participants

Infants were recruited at the maternity hospital in the city of XXXXXXX (XXXX). Families were informed about the study when babies were born. Those caregivers who expressed a willingness to be further informed were contacted when the babies were 6 months old (mo). The initial sample consisted of 143 infants born at term (more than 36 gestational weeks and weight over 2.7Kg) and did not present any family history of neurodevelopmental disorders (i.e., first and second-order relatives diagnosed with autism spectrum disorder (ASD) or attention deficit and hyperactivity disorder (ADHD)). Participants were followed up in a second session at the age of 9 months (n = 123), and a third session at the age of 16 to 18 months (n = 93). Most of the third sessions of data collection took place at the time of the COVID-19 pandemic. As requested by the health authorities of the country, the activity of the lab ceased for 4 months. As consequence, the age of the third session was extended from 16 to 18 months to facilitate the continuity of participants in the study. We will refer to it as the 16mo session hereon. The sample included varied according to the analysis required for testing each hypothesis. For instance, the trajectory changes included the three waves in the model, whereas stability analyses were done in pairs (e.g., 6mo session correlated to 9mo session). Further details can be found in Supplementary Tables 1 to 3 (see 2.4. Data Analysis).

### 2.2. Procedure

The rs-EEG protocol reported in this paper is part of a larger longitudinal study, which included other age-adapted protocols (eye-tracking tasks and different behavioral protocols at the 9mo and 16mo sessions) that were carried out before the EEG register. Data on other tasks and protocols will not be reported in this paper. Including all protocols, the approximated duration of the session was 30 minutes (6mo) or 1 hour (9mo and 16mo) including resting times between tasks, as well as setting up of the EEG acquisition net. The rs-EEG was the last protocol of the session and consisted of two 2-minutes blocks. In the first block, the experimenter blew soap bubbles in front of the baby, whereas the second block consisted of a video with geometrical shapes and soft music. During both periods of EEG acquisition, babies were seated on the parent’s lap and the parents were instructed to hold the baby comfortably and remain silent.

### 2.3. EEG

#### 2.3.1. EEG Register

We registered EEG brain activity using a high-density geodesic net (128 channels) and the software Net Station 4.3 (EGI Geodesic Sensor Net, Eugene, OR). The resting protocol was programmed and synchronized to Net Station with E – Prime 2.0. The signal was digitalized at 1000 Hz frequency, filtered using an elliptical lowpass (100 Hz) and high pass (0.1 Hz) hardware filters, and online reference to the vertex. The baby was also video-recorded in synchrony with the EEG acquisition. This video was used for posterior visual inspection to detect fragments in which the baby was fussy and/or there was a parental interruption (e.g., the parent signaling the video). Those fragments were marked as invalid and were rejected from the analysis in the segmentation step (see Anderson et al., 2022).

#### 2.3.2. EEG processing

The pre-processing of the EEG signal was conducted in the EEGlab (Delorme and Makeig, 2004) with the Maryland Analysis of developmental EEG (MADE) pipeline (Debnath et al., 2020). As the boundary electrodes (n = 20; see Supplementary Figure 1) were usually noisy, we removed them in the first step of the pre-processing. Then the continuous dataset was filtered with a Hamming window (0.2 – 48 Hz) using a finite impulse response filter (FIR). Bad channels were identified and removed with the FASTER plug-in (Nolan et al., 2010) and excluded until the interpolation step. After removing the channels, a copy of the dataset was performed and segmented into 1s segments. The dataset was copied, high-pass filtered (1 Hz), and cleaned from artifacts (threshold = ±1000 Hz). Then, ICA was performed to detect independent components containing blinks or ocular movements using infant adapted ADJUST plug-in (Leach et al., 2020). Those components were then copied into the original dataset and removed from the signal. Next, the original dataset was segmented into 2s epochs with a 50% of overlap. We applied a voltage threshold rejection (±125) to remove the rest of the artifacts of the signal. If any frontal channel exceeded the threshold, the segment was rejected as the frontal channels are more likely contaminated by residual artifacts of eye movements and blinks. In case, any other electrode surpassed the threshold, it was spherically interpolated. If more than 20% of channels were noisy in one segment, that segment was rejected. Then, the initially excluded channels by FASTER were recovered and spherically interpolated, and re-referenced to the average of all the channels. The remaining epochs were visually inspected and the segments with excessive noise were removed. Babies with less than 10 epochs (n = 13 at 6 mo.; n = 6 at 9mo; n = 2 at 16mo) or more than 10 electrodes interpolated were discarded from further analysis (n = 0).

#### 2.3.3. Absolute and relative power

Absolute power was computed employing the fieldtrip toolbox in MATLAB 2019a (Oostenveld et al., 2011) in each electrode with an FFT in 1 Hz steps from 1 to 45 Hz. The relative power was then computed by dividing the absolute power of a frequency band (e.g., alpha) by the power of all the ranges employed. We anchored the different frequency bands to individual alpha frequency (IAF; see below).

#### 2.3.4. Aperiodic and periodic components of the EEG signal

The signal was decomposed into periodic and aperiodic components with the FOOOF toolbox (Donoghue et al., 2020) called from fieldtrip. The FOOOF toolbox models the absolute power of the signal in each frequency (P(f)) as a combination of aperiodic (L(f)) and periodic (G; Σn Gn (f)) components. The aperiodic component is defined as L(f) = *b* – log[fx] where *b* is a constant offset and *χ* is the aperiodic exponent. The periodic contribution G is modeled as a gaussian peak. The parameters of the model were similar to previous studies (peak width limits: [2.5Hz – 12Hz], the maximum number of peaks: 5, aperiodic mode: fixed, peak threshold: 2; see Schawornkow and Voytek, 2021) and computed including the frequencies between 1 and 45 Hz. This range was larger than in previous studies with infants but necessary to compute the periodic brain activity in the faster frequency bands. Only participants with R^2^ > .95 were included in the model (Supplementary Table 4). Then, the aperiodic signal was computed and subtracted from the raw power to obtain the periodic part of the EEG activity.

#### 2.3.5. Frequency bands and clustering

As the frequency of each band changes along with development (Marshall et al., 2002), we cantered the frequency bands on the IAF. We considered IAF as the highest value between 5.5 and 9.5 Hz of the periodic power computed by the FOOOF toolbox (see Marshall et al., 2022, Stroganova et al., 1999) in the electrodes of the parietal cluster. If an infant did not present a clear IAF in any of the electrodes, we used a standard frequency band employed in infants (6 – 9 Hz). Then the frequency bands were constructed as follows: theta (0.6*IAF – 0.8*IAF), alpha (0.8*IAF – 1.2*IAF), beta (1.2*IAF – 20 Hz), and gamma (20 Hz – 45 Hz). As the main focus of this research was to compare the trajectories between periodic and relative power, we divided the periodic power into the same frequency bands that the relative power. We individually computed the values for each electrode and then averaged them over the parietal (Pz) and frontal (Fz) clusters (Supplementary Figure 1).

### 2.4. Data Analysis

#### 2.4.1. Change

As the attrition rate in developmental studies is high and missing data might bias the results, we employed maximum likelihood estimation in the analysis of children who came to at least two sessions (Enders, 2013; Graham, 2009; Matta et al., 2018). According to Littles’ test, missing data were missing completely at random (MCAR; *χ*2(7) = 4.80, p = .683). Also, the infants who performed the second and third sessions did not differ in socioeconomic status (SES) from those who did not come to the lab either in the second (t(91) = −.54, p =.59) or the third session (t(91) = −.41, p =.68), which indicated that the estimation would not be affected by relevant environmental factors. As a result, we decided to analyze the data employing linear mixed models estimating the missing data (n = 91; See Supplementary Table 1). The models included Age-Session (6, 9, 16months) and Cluster (Pz and Fz) as fixed repeated effects and participant as random effects for each of the frequency bands (theta, alpha, beta, and gamma) or aperiodic parameters (exponent and offset). We constructed the models following a top-down strategy (West, Welch & Galecki, 2006) and compared the final model with a chi-squared test for the godness of fit with AIC using the three common structures used in longitudinal studies (unstructured, compound symmetry, and autoregressive). In case model residuals were non-normally distributed, we employed the Tucker stair of ladders to transform the data and re-run the models from the first step. The degrees of freedom were computed using the Satterthwaite approximation. In case the fixed effects significantly interacted, we performed estimated means comparisons (EMs) corrected by Bonferroni.

Contrary to other parameters, we conducted a linear mixed model without estimating missing values (n = 23) for the alpha peak analysis because not all infants presented an alpha peak at 6mo and 9mo sessions. Therefore, we could not guarantee whether an infant who did not come to any of those sessions would or would not have had an alpha peak (Supplementary Table 2).

#### 2.4.2. Stability

To compute the stability between sessions, we ran two-tailed Spearman ranked correlations between each pair of sessions: 6mo – 9mo, 6mo – 9mo, and 9mo – 16mo without estimating the missing values (n = 72 in 6mo – 9mo correlation; n = 51 in 6mo – 16mo correlation; n = 28 in 9mo – 16mo correlation).

#### 2.4.3. Contributions of aperiodic and periodic EEG components to relative power

To determine the contribution of periodic and aperiodic components of the EEG to the relative power, we ran regression models for each frequency band and cluster. We included data from all sessions and introduced the Age-Session factor to account for developmental variability.

Therefore, the model had the relative power as a dependent variable (e.g., alpha parietal) and the session age, exponent, offset, and periodic power in the same frequency band and cluster as the predicted variable.

## 3. RESULTS

### 3.1. Change

#### 3.1.1. Aperiodic components

The offset increased over the sessions (main effect: *F* (12, 200.22) = 31.83, *p* < .001; EM: *p*s < .05), was larger in the parietal cluster (*F* (1, 191.945) =73.165, *p* < .001) and the interaction Age-Session × Cluster was significant (*F* (2, 278.20) = 6.48, *p* = .002; Fig. 1A). EMs unveiled a significant increase along the sessions in the frontal cluster (*p*s < .001) but only a significant augment between the first and the rest of the sessions in the parietal cluster (*p*s < .01). Also, the parietal cluster had a larger offset than the frontal cluster in the 6mo and 9mo sessions (*p*s < .001) but the difference was marginal at the 16mo session (*p* = .074).

**Figure 1.**
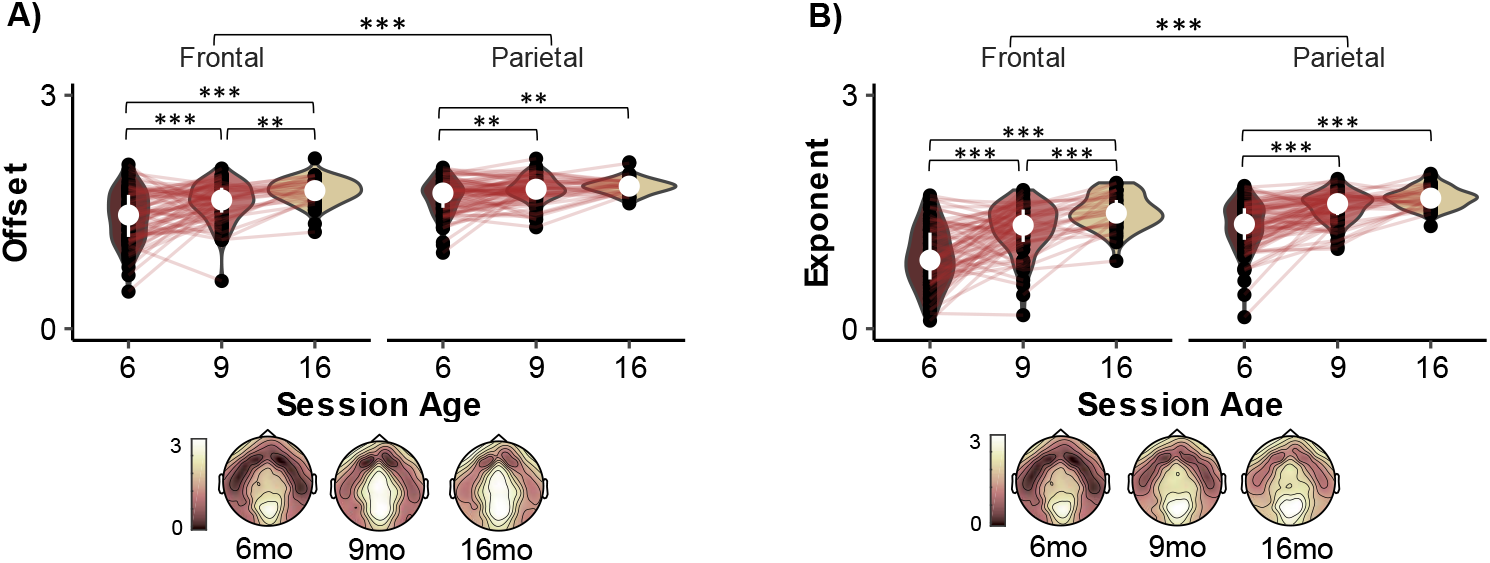
Aperiodic parameters development across sessions. Note. It displays a violin plot for each aperiodic component (offset and exponent) across sessions divided by cluster. Black dots represent individual values of each infant, whereas the white dot and white line correspond to the median and the first and third quartiles, respectively. The red lines that unite the black dots are the individual trajectory of infants who had two consecutive sessions. ** *p* < .01, *** *p* < .001.

The exponent component increased with age (Age-Session main effect: *F* (12, 211.01) = 79.22, *p* < .001; EM: *p*s < .001), displayed larger values in the parietal than the frontal cluster (*F* (1, 205.64) = 168.96, *p* < .001), and both factors significantly interacted (*F* (2, 278.33) = 7.52, *p* = .001; Fig. 1B). EMs revealed an increase from first to second and third sessions (*p*s < .001) but only marginal changes between second and third session in the parietal cluster (*p* = .054). In comparison, the exponent in the frontal cluster increased in all the waves (*p*s < .001). See Supplementary Table 5 for the descriptive values of the offset and exponent measures.

#### 3.1.2. Periodic brain activity

##### 3.1.2.1. Alpha peak

The percentage of infants with an alpha peak increased from 70.92% (6mo) to 89.01% (9mo) and 100% (16mo; Table 1 and Supplementary Table 6). Concerning the frequency of the alpha peak, we found a significant main effect of Age-Session (*F* (2, 44.32) = 30.11, *p* < .001; Fig. 2). There were no differences in the frequency of the alpha peak between the 6mo and 9mo Age-Sessions (*p* = 1) but the frequency of the alpha peak was slower at these two sessions compared to the 16mo session (*p*s < .001).

**Table 1.**
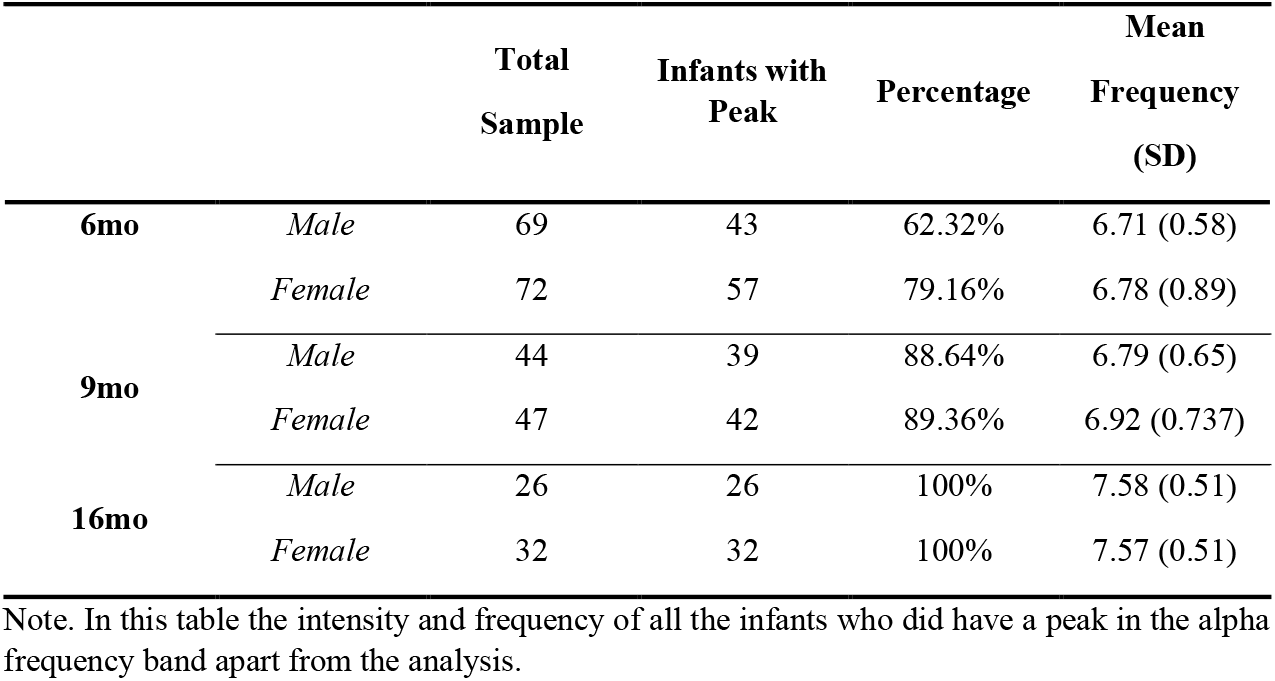
Percentage of infants with an Alpha peak and frequency (Hz) at the parietal cluster.

|  |  | **Total**<br>**Sample** | **Infants with**<br>**Peak** | **Percentage** | **Mean**<br>**Frequency**<br>**(SD)** |
| --- | --- | --- | --- | --- | --- |
| **6mo** | *Male* | 69 | 43 | 62.32% | 6.71 (0.58) |
|  | *Female* | 72 | 57 | 79.16% | 6.78 (0.89) |
| **9mo** | *Male* | 44 | 39 | 88.64% | 6.79 (0.65) |
|  | *Female* | 47 | 42 | 89.36% | 6.92 (0.737) |
| **16mo** | *Male* | 26 | 26 | 100% | 7.58 (0.51) |
|  | *Female* | 32 | 32 | 100% | 7.57 (0.51) |

Note. In this table the intensity and frequency of all the infants who did have a peak in the alpha frequency band apart from the analysis.

**Figure 2.**
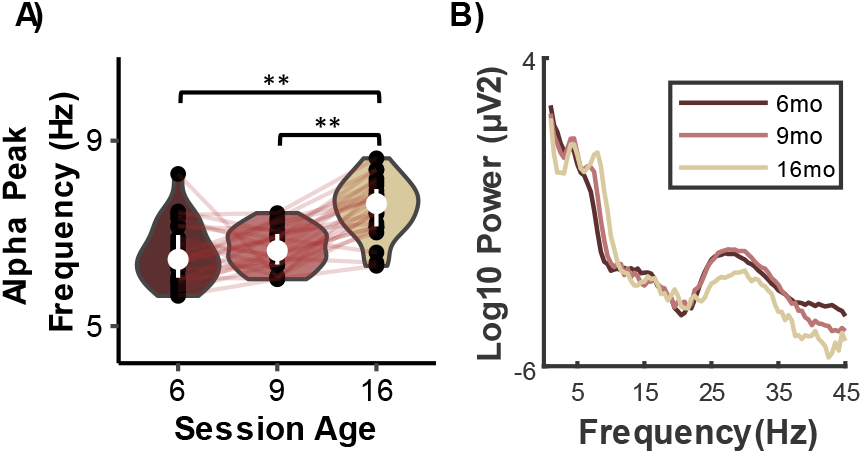
Alpha peak frequency development. Note. (A) Violin plot for the alpha peak frequency. Black dots represent the individual values of each infant, whereas the white dot and white line correspond to the median and the first and third quartiles, respectively. The red lines that unite the black dots are the individual trajectory of infants *** *p* < .001. The *p* values correspond to the EMs of the repeated measures ANOVA corrected by Bonferroni. (B) The power distribution of the periodic component from which the alpha peak has been computed.

##### 3.1.2.2. Theta periodic power

The parietal cluster displayed higher theta periodic power than the frontal one (*F*(1,260.82) = 6.01, *p* = .015), and it changed across Age-Session (*F*(2,264.91) = 11.86, *p* < .001). At 9mo, infants displayed greater theta power than at 6mo and 16mo (*p*s < .002), but there were no differences between the 6mo and the 9mo sessions (*p* = 1; Fig 3A and Supplementary Table 7 for the descriptive values of periodic power of all frequency bands).

**Figure 3.**
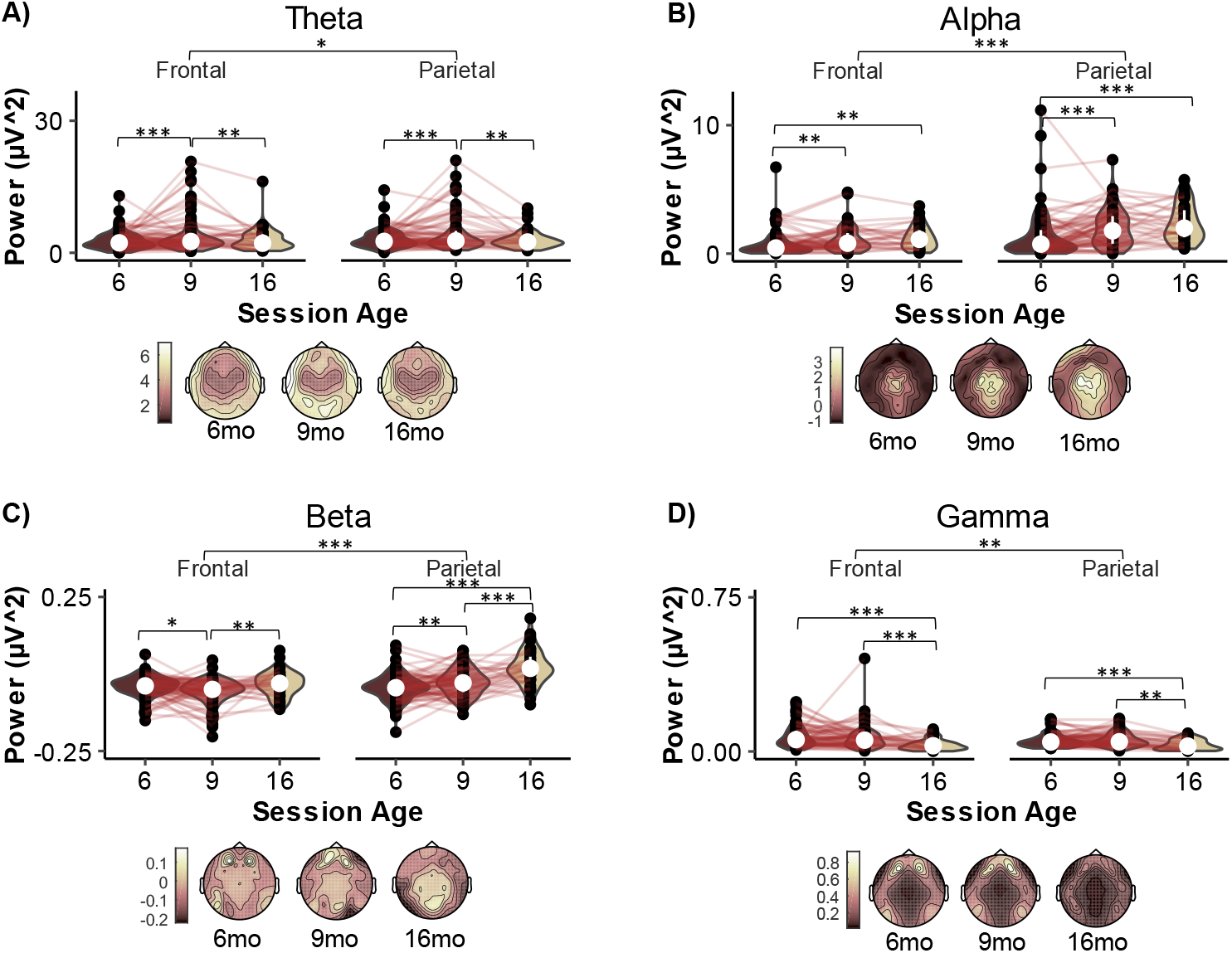
Periodic Power Development. Note. The figure displays a violin plot of the periodic power for each frequency broadband, Theta (A), Alpha (B), Beta (C) and Gamma (D), separated by cluster (Parietal and Frontal). Black dots represent the individual values of each infant connected across time by a red line. The white dot and lines represent the median and the first and third quartiles, respectively. Below the figure shows the topographical map of the relative power in each session. *** *p* < .001, ** *p* < .01, \**p* < .05. The *p* values correspond to the EMs of the LMM corrected by Bonferroni.

##### 3.1.2.3. Alpha periodic power

We found higher alpha power at the parietal cluster than the frontal cluster (*F*(2,241.46) = 178.46, *p* < .001). Also, we obtained a significant main effect of Age-Session (*F*(2,212.43) = 22.405, *p* < .001; EMs: *p*s <.001; Fig 3B), although we did not find significant differences between 9mo and 16mo of age (EM: *p* = 1). The Age-Session x Cluster interaction was also significant (*F*(2,282.11) = 3.71, *p* = .026). There was an increase of alpha power from the first to the second/third sessions (*p*s < .009) but without change between the 9mo and 16mo sessions (*p*s > .689) in the frontal and parietal clusters. Moreover, the parietal cluster showed greater alpha power than the frontal cluster in all the sessions (*p*s < .001).

##### 3.1.2.4. Beta periodic power

The periodic power in the beta band was higher at the parietal than the frontal cluster (*F* (1, 182.02) = 60.93, *p* < .001) and varied between sessions (*F* (2, 194.63) = 24.97, *p* < .001; Fig 3C). Beta power did not change between 6 and 9 months of age (*p* = 1) but increased at the 16mo session (*p*s < .001). The Age-Session × Cluster interaction (*F* (2, 281.50) = 20.67, *p* < .001), and subsequent EM revealed an increase in periodic power between 6mo and 9mo (*p* = .007) and between 9mo and 16mo (*p* < .001) infants at the parietal cluster. At the frontal cluster, beta power decreased from 6mo to 9mo (*p* = .021) and then it increased from 9mo to 16mo (*p* = .001). Furthermore, both clusters had equal power in the first session (*p* = .629), but power was higher at the parietal cluster in the 9mo and 16mo sessions (*p*s < .001).

##### 3.1.2.5. Gamma periodic power

The periodic power in gamma frequency band was larger in the frontal cluster (*F* (1,168.94) = 8.77, *p* = .004) and changed with age (*F*(2,195.97) = 15.81, *p* < .001; Fig. 3D), not showing an interaction with cluster (*F* (2, 284.28) = 2.12, *p* = .121). The power at 6mo and 9mo was higher than the power at 16mo (*p*s < .001), with no difference between the first two sessions (*p* = 1).

#### 3.1.3. Relative power

##### 3.1.3.1. Theta relative power

Relative power in theta was larger in the frontal cluster (*F*(1,229.70) = 72.44, *p* < .001), decreased throughout Age-Session (main effect: *F*(2,208.31) = 13,83, *p* < .001; EMs: *p*s < .005; Fig 4A), and showed a significant Session x Cluster interaction (*F*(2,277.22) = 5.83, *p* = .003). In the frontal cluster, there was a decrease only between the first and the third session (*p* = .01) but not in the other pairwise comparisons (*p*s > .125). On the contrary, theta relative power showed a significant decrease at the parietal cluster in all Age-Sessions (*p*s > .002). See Supplementary Table 7 for the descriptive information on the relative power for all the frequency bands.

**Figure 4.**
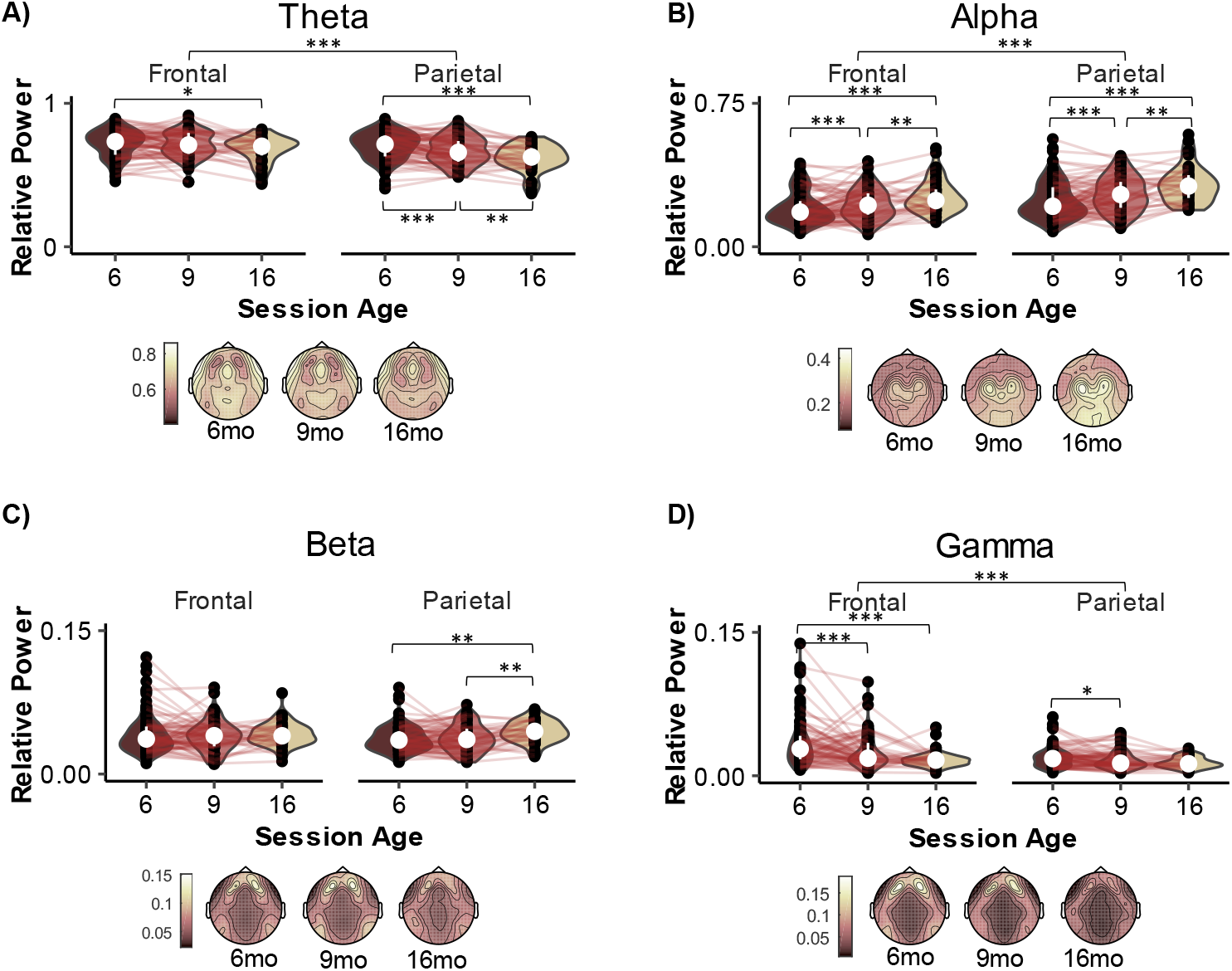
Relative Power Development. Note. The figure displays a violin plot of the relative power for each frequency broadband, Theta (A), Alpha (B), Beta (C) and Gamma (D), separated by cluster (Parietal and Frontal). Black dots represent the individual values of each infant connected across time by a red line. The white dot and lines represent the median and the first and third quartiles, respectively. Below the figure shows the topographical map of the relative power in each session. *** *p* < .001, ** *p* < .01, \**p* < .05. The *p* values correspond to the EMs of the LMM corrected by Bonferroni.

##### 3.1.3.2. Alpha relative power

Relative power in alpha increased throughout Age-Session (main effect: *F*(2,255.05) = 19.57, *p* < .001; EMs: *p*s < .01; Fig 4B) and was higher in the parietal cluster than in the frontal cluster (*F*(1,255.71) = 138.04, *p* < .001).

##### 3.1.3.3. Beta relative power

We obtained a main effect of Age-Session (*F*(2,239.11) = 5.43, *p* = .005; Fig 4C) for beta relative power, which only significantly changed between the first and second to the third session (*p*s < .033). There were no differences between the first and second sessions (*p* = 1). The Session × Cluster interaction was significant (*F*(2,284.47) = 4.41, *p* = .005). The frontal cluster did not vary across Age-Sessions (*p*s > .161), whereas relative beta power in the parietal cluster increased between 6mo and 9mo in comparison to 16mo (*p*s < .003). At 6mo, babies had less beta power at the parietal cluster compared to the frontal cluster (*p =* .003), but these differences were not significant at 9 or 16 months of age (*p*s > .182).

##### 3.1.3.4. Gamma relative power

Gamma relative power was larger in the frontal cluster (*F*(1,224.85) = 89.36, *p* < .001) and decreased (*F*(2,223.86) = 14.88, *p* < .001; Fig. 4D) between 6 and 9 or 16 mo (*p*s < .001), showing no differences between 9 and 16 month-old participants (*p* = .250). The Age-Session × Cluster interaction was significant (*F*(2,284.33) = 4.67, *p* = .01) and revealed different trajectories for each cluster. Gamma relative power decreased between the 6 and 9 or 16-month-old sessions (*p*s <. 001) at the frontal cluster, whereas we only found a significant reduction between 6mo and 9mo (*p* = .041). No other comparison where significant (*p*s > .064).

### 3.2. Within-subjects stability of the measures

#### 3.2.1. Aperiodic components

At the parietal cluster, both aperiodic components were correlated between the first and the rest of the sessions (r_s_ = [.33 .39], *p*s < .01), although between 9 and 16 month-old sessions the correlation was only significant for the offset (r_s_ = .45, *p* < .001). In the frontal cluster, we found a positive correlation for offset between the 6mo and the 16mo sessions (r_s_ = .454, *p* < .01) while both slope (r_s_ = .36, *p* < .01) and offset (r_s_ = .36, *p* < .001) were positively correlated between 9 and 16 month-old infants (see Table 2).

**Table 2.** Stability of the aperiodic parameters (slope and offset) and periodic power between sessions

|  |  | 9-month-old Session |  |  |  |  |  | 16month-old Session |  |  |  |  |  |
| --- | --- | --- | --- | --- | --- | --- | --- | --- | --- | --- | --- | --- | --- |
|  |  | *Aperiodic* |  | *Periodic Power* |  |  |  | *Aperiodic* |  | *Periodic Power* |  |  |  |
|  | Session | Slope | Offset | Theta | Alpha | Beta | Gamma | Slope | Offset | Theta | Alpha | Beta | Gamma |
| **Parietal** | *6mo* | .35** | .36*** | .60*** | .46*** | .26* | .38*** | .49*** | .546*** | .61*** | .52*** | .26 | .59*** |
|  | *9mo* | N/A | N/A | N/A | N/A | N/A | N/A | .13 | .45** | .66*** | .57** | .49** | .54** |
| **Frontal** | *6mo* | .22 | .22 | .51*** | .28* | .24* | .39*** | .30* | .38* | .52** | .61*** | -0.04 | .196 |
|  | *9mo* | N/A | N/A | N/A | N/A | N/A | N/A | .46** | .42* | .59*** | .51** | .16 | .19 |

Note. \* $p < .05$ ; \*\* $p < .01$ ; \*\*\* $p < .001$

#### 3.2.2. Periodic brain activity

Values of the frequency of the alpha peak were correlated between the first and second Age-Session (r_s_ = .34, *p* < .05) and between the second and third Age-Session session (r_s_ = .209, *p* < .05), whereas the 6mo to 16mo correlation did not reach a significant level (r_s_ = .21, *p* > .05).

At the parietal cluster, the 6mo periodic power was positively correlated with that of the 9mo and 16mo sessions in all the frequency bands (r_s_ = [.26 .61], *p*s < .05), except for beta at 16mo, which was only marginal(r_s_=.26, *p* = 0.074). Also, the periodic power at 9mo was related to 16mo periodic power in all the frequency bands (r_s_=[.49 .66], *p*s < 0.01). Similarly, at the frontal cluster the power at 6mo was correlated to that at 9mo and 16mo in all frequency bands (r_s_ = [.24 .61], *p*s < .05) with the exception of beta (r_s_=-.04, *p* = .775) and gamma (r_s_ = .196, *p* = .187) bands at 16 mo. The periodic power at 9mo was positively correlated with 6mo power in theta (r_s_ = .59, *p* < .001) and alpha (r_s_=.51, *p* < .001) but no other frequency bands ((r_s_ = [.16 .19], *p*s < .297); Table 2).

#### 3.2.3. Relative power

The relative power at the parietal cluster at 6mo was positively correlated to the rest of Age-Sessions in all the frequency bands, (r_s_ = [.38 .60], *p*s < .01), which also occurred between the 9mo and 16mo sessions (r_s_ = [.54 .67], *p* < .001). At the frontal cluster, 6mo relative power was positively correlated in all frequency bands (r_s_ =[.29 54], *p*s < 0.01) but gamma (r_s_=.20, *p* = .071) at 9mo, and it was also related to 16mo relative power in all the frequency bands (r_s_= [.30 .61], *p* < .05). The relative power between 9 and 16 month-old correlated in all the frequency bands(r_s_= [.48 .61], *p*s < .001) with the exception of gamma frequency band (r_s_=.22, *p* = .218; Table 3).

**Table 3.**
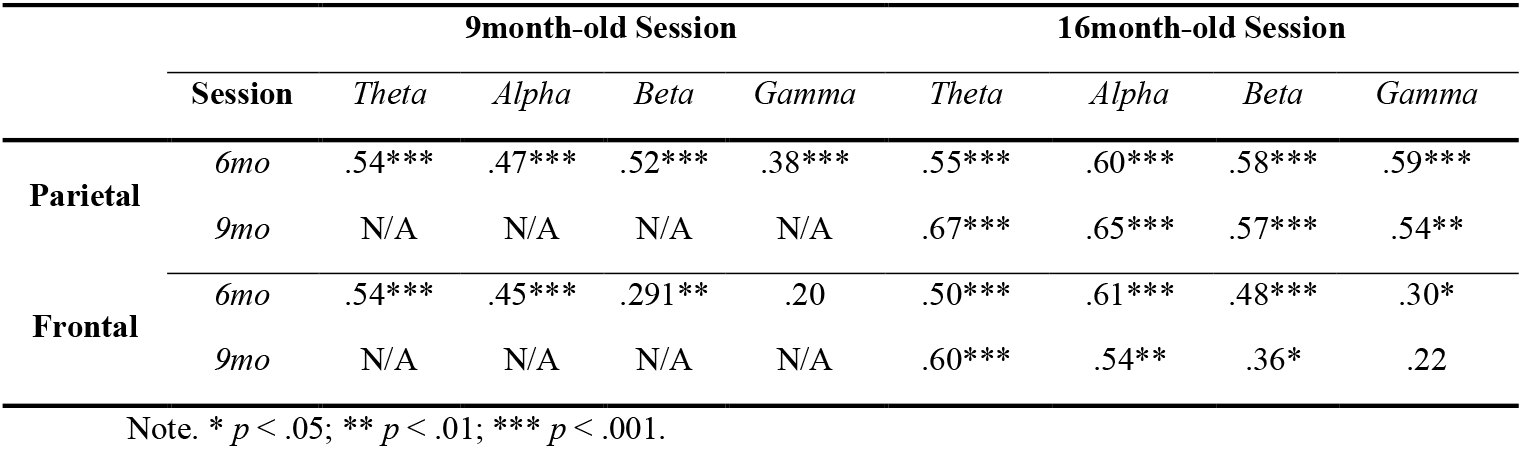
Stability of the relative power between sessions

|  |  | 9month-old Session |  |  |  | 16month-old Session |  |  |  |
| --- | --- | --- | --- | --- | --- | --- | --- | --- | --- |
|  | Session | Theta | Alpha | Beta | Gamma | Theta | Alpha | Beta | Gamma |
| **Parietal** | 6mo | .54*** | .47*** | .52*** | .38*** | .55*** | .60*** | .58*** | .59*** |
|  | 9mo | N/A | N/A | N/A | N/A | .67*** | .65*** | .57*** | .54** |
| **Frontal** | 6mo | .54*** | .45*** | .291** | .20 | .50*** | .61*** | .48*** | .30* |
|  | 9mo | N/A | N/A | N/A | N/A | .60*** | .54** | .36* | .22 |

Note. \* $p < .05$ ; \*\* $p < .01$ ; \*\*\* $p < .001$ .

### 3.3. Contribution of aperiodic and periodic activity to relative power

We carried out regression models to understand the contribution of periodic power and aperiodic components of the EEG to the relative power in all frequency bands. See Table 4 for the complete information about the regression models.

**Table 4.**
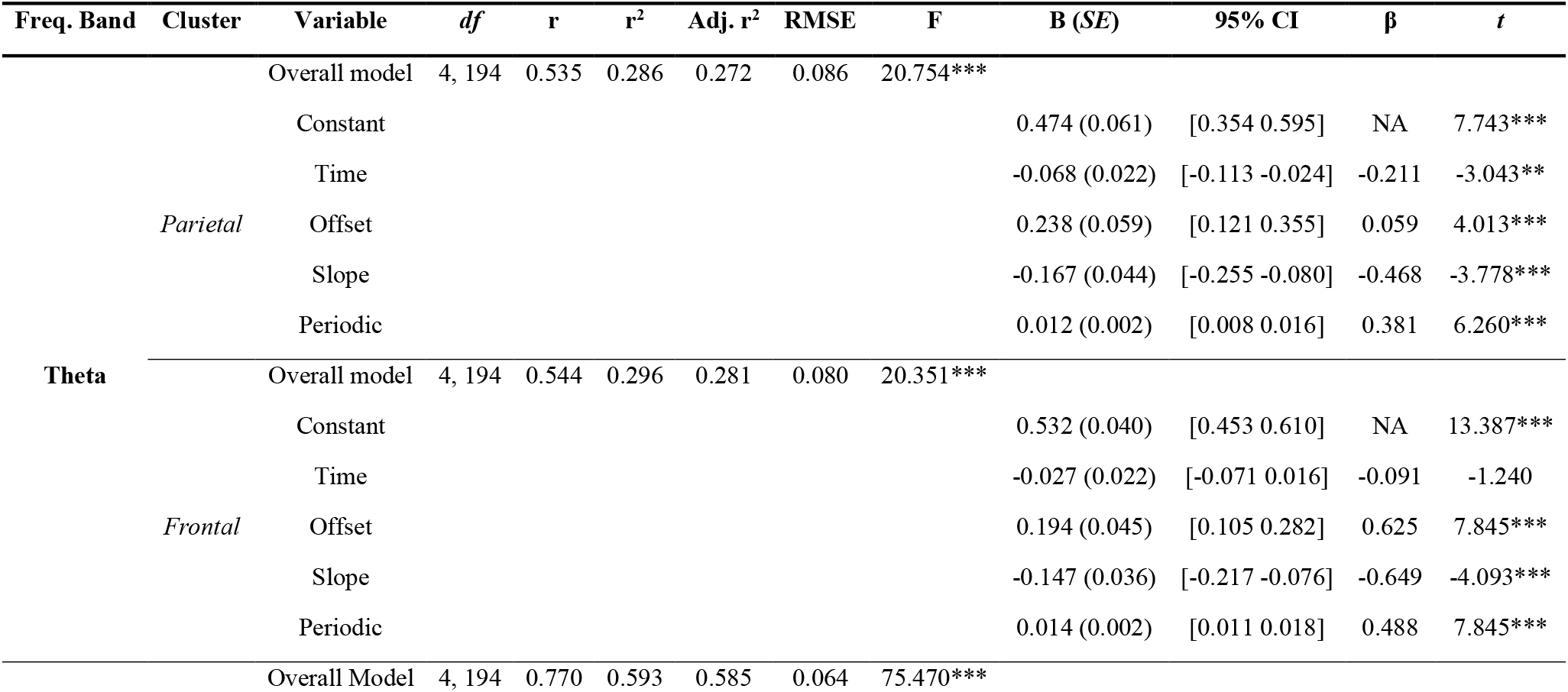
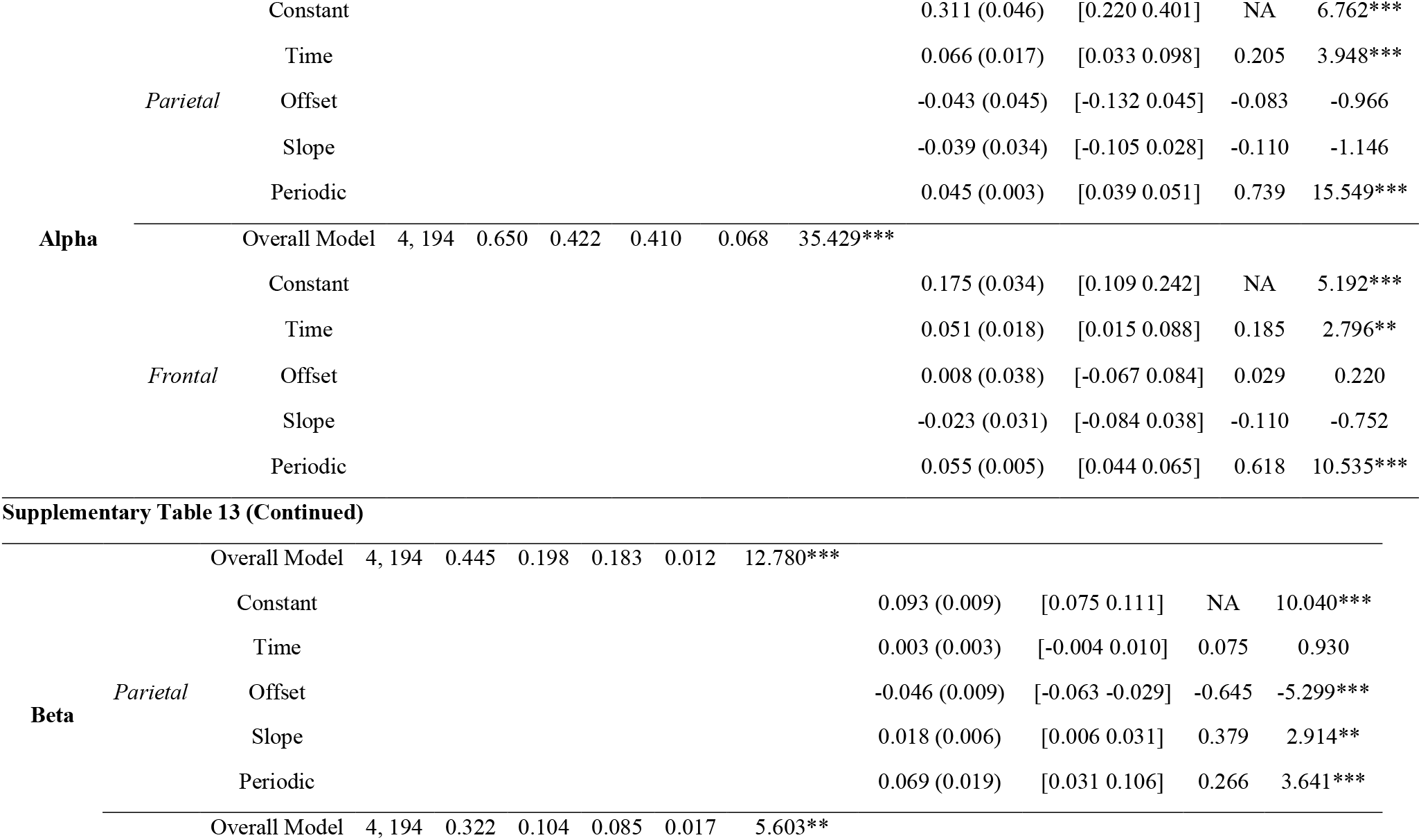
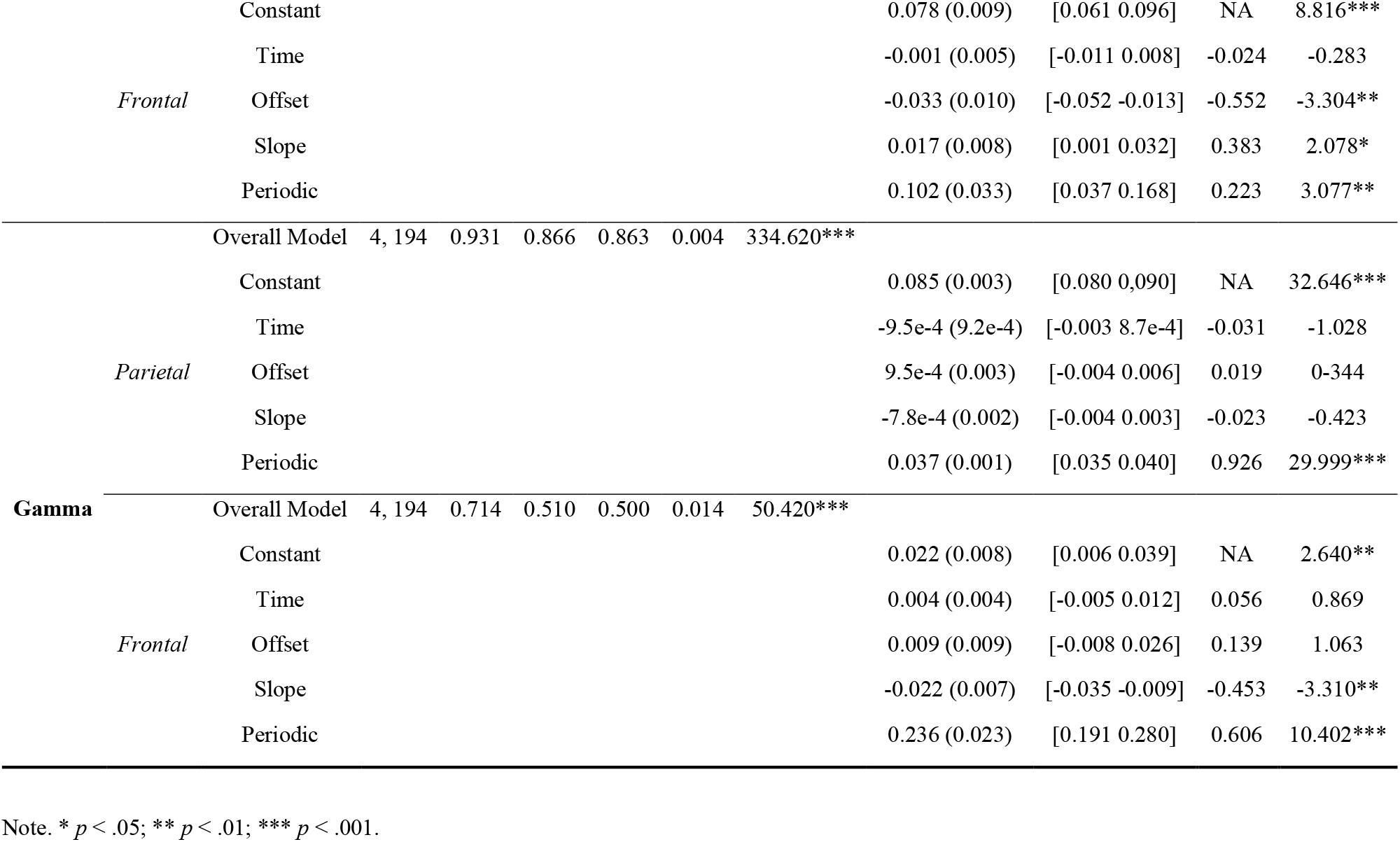
Regression between relative power and the aperiodic components and periodic power for each frequency band and cluster. It displays the general parameters of the model and the predictor parameters.

#### 3.3.1. Theta

Regarding to theta band, the regression models were significant for both clusters (parietal: adj. R^2^ = .27, F (4,194) = 20.75, p < .001; frontal: adj. R^2^ = .28, F (4,194) = 20.35, p < .001). In the parietal cluster, relative power was positively predicted by offset (std. β = .0.06, p < .001) and periodic power (std. β = .38, p < .001). On the contrary, it was negatively predicted by time (std. β = −.21, p < .001) and slope (std. β = −.47, p < .001). Similarly, the relative power in the frontal cluster was positively related to offset (std. β = .64, p < .001) and periodic power (std. β = .49, p < .001), and negatively to slope (std. β = −.65, p < .001). However, it was not related to time (std. β = −.091, p > .05).

#### 3.3.2. Alpha

The regression model in alpha relative power was significant at the parietal (adj. R^2^ = .59, *F* (4, 194) = 75.47, *p* < .001) and frontal (adj. R^2^ = .41, *F*(4,194) = 35.43, *p* < .001) clusters. In both clusters, time (parietal: std. β = .21, *p* < .001; frontal: std. β = .19, *p* < .001) and periodic power (parietal: std. β = .74, *p* < .001; frontal: std. β = .62, *p* < .001) were positively related to relative power. Any of the aperiodic parameters were significant contributors to alpha relative power.

#### 3.3.3. Beta

Beta relative power was significantly predicted in both clusters (parietal: adj. R^2^ = .18, *F* (4,194) = 12.79, *p* < .001; frontal: adj. R^2^ = .09, *F*(4,194) = 5.60, *p* < .01). The significant predictors for the parietal and frontal clusters were identical: offset (parietal: std. β = −.65, *p* < .001; frontal: std. β = −.552, *p* < .01), slope (parietal: std. β = .38, *p* < .01; frontal: std. β = .38, *p* < .05), and periodic power (parietal: std. β = .266, *p* < .001; frontal: std. β = .22, *p* < .01). Age-Session was not a significant predictor for any of the clusters (*p*s > .05).

#### 3.3.4. Gamma

Regression models yielded significant R^2^ for relative gamma power at both frontal and parietal clusters (parietal: adj. R^2^ = .86, *F* (4,194) = 334.62, *p* < .001; frontal: adj. R^2^ = .5, *F* (4,194) = 50.42, *p* < .001). At the parietal level, the gamma relative power was only predicted by the periodic power (std. β = .93, *p* < .001), whereas in the frontal cluster the model included both slope (std. β = −.45, *p* < .01) and periodic power (std. β = .61, *p* < .001) as significant predictors.

## 4. DISCUSSION

The goal of the present study was to characterize the trajectory of relative power, periodic power, and aperiodic components and test its stability in a baseline rs-EEG in infants. Also, we aimed for disentangling the contributions of periodic power and aperiodic background to relative power. Our results were similar and expanded the findings in relative power (e.g., reduction of theta relative power) and suggest differential developmental curves when the power is isolated from the aperiodic background. Additionally, the measures were stable across sessions, especially in the parietal area. Furthermore, the relative power was related to the aperiodic components in theta, beta, and gamma frequency bands but not in the alpha frequency band, which suggest that EEG relative power captures a combination of both periodic and aperiodic brain activity.

### 4.1. EEG power: trajectory, stability, and relationship

#### 4.1.1. Aperiodic Components

The exponent and offset components of the aperiodic signal increased with age in our study, which contrasts with previous studies in infants and children and our initial hypothesis (e.g., Cellier et al., 2021; Schaworonkow, & Voytek, 2021; Hill et al., 2022; Voytek, 2015; Donoghue et al., 2020a; McSweeney et al., 2021). However, the measures were stable, and the exponent and the offset at the parietal cluster were good predictors of infant values in later sessions. Conversely, the frontal cluster displayed less stability, which might be related to the rapid maturation of the frontal lobe in this period (Bell & Fox, 1992).

Our results indicate that the aperiodic background curve becomes steeper between six months and 18 months of age, decaying faster and starting with higher energy when the frequency range is wider. This would suggest a change toward more inhibitory activity in the excitatory/inhibitory balance (Gao, Petersen &, Voytek., 2017; Voytek & Knight, 2015). However, it is also likely that differences between studies might account for this variability.

Concerning the protocol variances between studies, the only experiment with infants to date (Schaworonkow, & Voytek, 2021) could not isolate moments without baby-caregiver interaction, which probably impacted the EEG measures (Johnson et al., 2016). Also, in comparison with the protocols for children and adults (eyes open/closed protocol), our study presented external stimuli (soap bubbles and dynamic patterns of color and shapes in combination with soft music) to calm the infants and reduce their movement (Saby & Marshall, 2012; see also Anderson, Perone, & Garstein, 2022). Consequently, the presence of stimuli could have impacted the rs-EEG power pattern, as our protocol involves the baby watching the bubbles or the video. Indeed, a study by Hill and colleagues (2022) found different developmental trajectories between eyes closed vs. eyes open condition after. Also, some authors have focused on one of the conditions (e.g., Cillier et al., 2021). In addition, a recent combined rs-fMRI/EEG study by Jacob and colleagues (2021) in adults found that the aperiodic parameters were related to the hemodynamic changes in the auditory-salience network, which reflect the likely impact of the external stimuli in electrophysiological measures. Thus, our baseline rs-EEG might have been more influenced by external stimulation, whereas other rs-EEG protocols with older children and adults might be more related to a mind-wandering state.

Another possible influence that might have caused differences in the results between previous studies and ours is the pre-processing steps. In infant studies, authors have employed a narrower range of frequencies to compute the aperiodic parameters. For instance, Schaworonkow and Voytek (2021) analyzed the 1 to 10 Hz range, and Shuffrey and colleagues (2022) divided the range into 1 to 20 Hz and 21 to 41 Hz. These changes might affect the aperiodic curve, especially in infants with more motor artifacts that affect beta and gamma, frequency bands. To test this idea, we re-computed the aperiodic parameters with a 1 to 10 Hz range and the pre-processing parameters proposed by Schaworonkow and Voytek (2021). In that frequency range, we found a reduction in the exponent and an increase in the offset between the 6mo and the following sessions (see Supplementary Results and Supplementary Fig. 3 and 4). Thus, a wider range of frequencies in the parametrization of the aperiodic components affects the curve slope trajectory to fit higher frequencies. Indeed, the study of Shuffrey and colleagues (2022) found that the exponent between the 1 to 20 Hz range was larger than the exponent of the 21 to 40 Hz range in new-borns, which signals a steeper reduction of the aperiodic curve in the lower frequencies, which might be related to the presence of noise. Therefore, dividing the curve into low-frequency and high-frequency or computing the aperiodic parameters with a knee would be probably beneficial for the fit of the aperiodic background curve in infants. However, in studies with children and adolescents who can follow explicit instructions, the power-spectrum curve is less affected by motor artifacts which might facilitate the replicability even employing a wider frequency range.

#### 4.1.2. Periodic and relative power

The development of relative compared to periodic power varied in two of four frequency bands. The relative power in theta diminished between 6 and 18 month-old at the parietal cluster, whereas the periodic power displayed an inverted “u” shape trajectory. Similarly, beta relative power increased at the parietal cluster in the age range of our study, but the periodic power increased in this area from 6 to 9 months of age and then decreased. Alpha and gamma frequency bands showed the same pattern of age-related change for both relative and periodic power, augmenting and decreasing, respectively. Additionally, infants who presented an alpha peak increased a 30%, from 70% to 100% in the age range studied. Also, the mean individual alpha frequency peak increased by nearly 1Hz.

The developmental trajectories in relative power were similar to previous experiments with infants: reduction in theta relative power and increment of alpha relative power (Marshall et al., 2002; Stroganova et al., 1999; Orekhova et al., 2006). The maturation curve found in gamma had the same direction as the experiment by Tierney and colleagues (2012). Only the beta band development differed from previous studies as relative power increased in our study (Tierney et al., 2012). This difference might be because we anchored beta to IAF instead of choosing a predefined band, which likely affected the computation of the relative power.

The observed changes in periodic power fit previous literature to some extent. First, the increase in alpha peak frequency is similar to the findings of Marshall and colleagues (2002). Indeed, the alpha peak found after removing the background activity remains in the range called “infant alpha” (6 - 9 Hz; Stroganova et al., 1999; Orekhova et al., 1999, 2001, 2006), and it will probably transit to adult frequencies around the seventh year of life (Cellier et al., 2021). Additionally, we observed an increment of periodic power in alpha, which is consonant with Schaworonkow and Voytek’s (2021) results in younger infants (increment of an alpha burst presence over the first seven months of life). With high-frequency bands, both beta and gamma showed changes in this period, reflecting a rapid reconfiguration of brain activity above the aperiodic curve.

Electroencephalography power measures were relatively stable in the transition between infancy and toddlerhood. That is to say, the relative power in previous sessions was a predictor of the follow-up sessions in all the frequencies and clusters, except for frontal gamma. On the other hand, periodic power in earlier sessions is predicted to a lesser extent than the relative power of future power. For instance, beta periodic power only was correlated between the 6mo and 9mo sessions, and gamma displayed less stability than the rest of the frequency bands. These results are consonant with the stability alpha relative power between 10mo and 14mo and 14mo to 24mo, but not between 5 and 10 months of age infants, found by Marshall and colleagues (2002). The differences between the correlation between 6mo and 16mo might be due to the change from 5 to 6 month-old, which would suggest that the EEG becomes more stable in this period. This is also in consonance, in combination with aperiodic stability results, with a recent study by Demuru & Fraschini (2020) that showed the sensibility of aperiodic parameters to individual variations in adults. Given this, rs-EEG measures might be considered a fingerprint of the individual brain activity, combining both change and stability across the development. However, the measure selected seems to affect stability. In our study, relative power significant correlations between sessions were more present than in the periodic power. This might point out a constant ratio between frequencies along with development. Despite that, it can also be related to how we computed the periodic power (i.e., power above the aperiodic background curve in canonical bands), and employing a more individual parametrization might improve the stability (see 4.3. Limitations).

Finally, and more importantly, we have shown that the relative power is a mixture of both aperiodic and periodic components in most of the frequency bands. For instance, theta and beta relative power was predicted by the offset and exponent parameters. Also, frontal gamma included the exponent as a significant predictor. Only alpha (frontal and parietal) and gamma (parietal) were explained by periodic power without the contribution of other aperiodic components of the signal. However, it does not imply that alpha and gamma are not related to exponent and offset, but that the periodic power predicts the variability to a greater extent to exclude the aperiodic parameters of the model. Therefore, relative power captures to some extent the underlying oscillations above the aperiodic background activity, but it is usually a mixture of the background activity and the periodic power. Our results are consonant with previous research conducted by Donoghue and colleagues (2020a) in a longitudinal dataset from childhood to older-adults populations (Multimodal Resource for Studying Information Processing in the Developing Brain). In their study, they compared the correlation between aperiodic parameters and the periodic power to several power ratios and found that alpha ratios were more related to the direct power of alpha than the aperiodic parameters, but the opposite pattern emerged when the ratios did not include the alpha frequency band (e.g., theta/beta). Altogether shows the relevance of considering the aperiodic components when working with EEG power. Furthermore, it indicates that alpha power might be biasing the results of the rest of the frequency bands when computed relative power. That is to say, the large increment in alpha periodic activity observed in early development probably overshadows age-related changes in other frequency bands that do not present a clear peak.

### 4.2. EEG power development and neurodevelopmental disorders

As the relative power contains combines aperiodic and periodic brain activity, previous brain-behavior relationships can be driven by any of them, which could have impacted the replicability between studies (Donoghue et al., 2020a, 2020b Voytek & Knight, 2015). Thus, separating the EEG signal into periodic and periodic components provides a way to test separately their impact.

One example of a well-studied phenomenon is the relationship between the theta/beta ratio and ADHD (Ogrim, Kropotov, & Hestad, 2012; Liechti et al., 2013). Recent studies showed that a higher theta/beta ratio was related to a higher risk of ADHD in 10mo infants (Begum-Ali and colleagues, 2022). However, the study by Donoghue and colleagues (2020a) found that aperiodic components conflate the theta/beta ratio. Furthermore, recent research by Karalunas and colleagues (2021) found that 1mo infants with ADHD risk had a steeper aperiodic background. More interestingly, the relationship between aperiodic components and ADHD in adolescents was sensitive to whether the participant had taken medication (Ostlun et al., 2021, Robertson et al., 2019).

Another of the most studied relationship of relative power concern ASD. However, to date, the findings are contradictory sometimes and malleable with development. For instance, a review published in 2013 points out an augment of low-frequencies power, beta, and gamma, but a reduction in alpha power in the population diagnosed or with risk of ASD (Wang et al., 2013). However, other studies in infants signal that the trajectory of relative power is different in those at risk of ASD with less power in beta and gamma (Tierney et al., 2012). Additionally, Gabard-Durnam and colleagues (2019) found that lower gamma power was consistently related to ASD risk in infancy and toddlerhood. However, a recent study by Shuffrey and colleagues (2022) found that infants’ EEG aperiodic slope at birth was related to later ASD symptoms, whereas absolute power was not associated with ASD symptomatology.

### 4.3. Limitations

The current study focused on the early development of relative power in different frequency bands and the possible contribution of aperiodic and periodic in the canonical frequency bands up to gamma (3Hz approx. to 45Hz). Thus, our range is wider than most infant studies employing rs-EEG and might have impacted the signal quality because high-frequency bands are easily affected by motor artifacts and eye movements. However, we took particular care in processing the EEG signal to minimize the impact of movements. For instance, we visually inspected each participant’s video and marked motor artifacts, which were removed from the signal. Also, we employed ICA to remove blinks and eye movement and eliminated the segments with frontal activity over the threshold. Nevertheless, in case the residual gamma was excessive due to the movements, it would be expected to increase the power of gamma across age, as the blinks and eye movements increase from 6 to 16 month-old.

Concerning pre-processing, we wanted to compare the periodic oscillations and relative power in a similar way to traditional approaches. Therefore, we decided to compute periodic power as we did in the relative power: anchoring the frequency bands to IAF in canonical ranges. Further studies would benefit from finding each frequency peak or employing more advanced techniques to determine the individual range of each frequency (Cohen, 2021; Ostlun et al., 2022). Also, especially in infancy, oscillatory brain activity appears in bursts and is not constant across the entire register (Jones, 2016). Therefore, testing the appearance of those bursts and computing the power with them would probably increase the reliability of results and give other parameters of interest (Cole & Voytek, 2019; Ostlun et al., 2022; Rayson, et al., 2022). Finally, as infants protocols incorporate an active baseline, registering peripheral and behavior measures related to infants’ cognitive state could give us information and increase the comparability between infant and other developmental studies (Orekhova, Stroganova, & Posikera, 2001; Xie, Mallin, & Richards, 2018).

### 4.4. Conclusion

Relative power is the most common analysis approach to studying rs-EEG signals. Patterns of the relative power of different frequency bands show fast changes in the early years of life and are associated with cognitive development and neurodevelopmental disorders risk. Although it is usually assumed that relative power represents neural oscillations, the EEG signal combines both aperiodic and periodic components. Our results indicate that, at least in the transition from infancy to toddlerhood, changes in relative power are partially driven by changes in aperiodic activity. Therefore, relative power seems to capture both aperiodic and periodic components of the EEG power instead of the putative oscillations of EEG. Also, studies with neurodevelopmental disorders suggest that aperiodic background is affected in infants at risk. This signal that relative power might not be sensitive enough alone and it the necessity to incorporate more fine-grained measurements of EEG power to unveil the mechanism underlying brain maturation and its relation to cognitive processes and brain disorders.

## Supporting information

Supplementary_Results

## Acknowledgments

We would like to thank all the families that took part in this study. The research presented in this paper was funded by the National Research Agency of Spain (PSI2017-82670-P) awarded to M.R.R., and a pre-doctoral fellowship in neuroscience was granted to JRP by the Tatiana PGB foundation.

