## Supplementary_Tables_and_Figures for "Early development of electrophysiological activity: contribution of periodic and aperiodic components of the EEG signal"

**SUPLEMENTARY MATERIAL**

### Supplementary Table 1

*Demographical information of the sample was included in the linear mixed models' analysis. The table represents the mean (SD) of the gestational weeks, gestational weight, and the age of the participants in each one of the sessions.*

| | | Gestation | | Session Age (days) | | |
|---|---|---|---|---|---|---|
| | *n* | *Weeks* | *Weight (grams)* | *1st Session* | *2nd Session* | *3rd Session* |
| **Male** | 40 | 39.49 (1.47) | 3417.37 (541.69) | 193.75 (9) | 283.12 (8.1) | 520.74 (28.19) |
| **Female** | 51 | 39.48 (1.34) | 3311.34 (396.94) | 194.71 (9.05) | 285.89 (10.96) | 518.45 (21.52) |
| **Total** | 91 | 39.49 (1.39) | 3358.74 (467.27) | 194.28 (8.99) | 284.63 (9.81) | 519.37 (24.25) |

Note. The gestational period, weight, and the age of the first session were computed with the 91 infants included in the analysis of linear mixed models. The age of the second and third sessions was computed from those infants who came to the laboratory, 76 and 44, respectively.

### Supplementary Table 2

*Demographical information of the infants included in the alpha peak linear mixed model. It represents the mean (SD) of the gestational weeks, gestational weight, and the age of the participants in each one of the sessions.*

| | | Gestation | | Session Age (days) | | |
|---|---|---|---|---|---|---|
| | *n* | *Weeks* | *Weight (grams)* | *1st Session* | *2nd Session* | *3rd Session* |
| **Female** | 8 | 38.52 (1.99) | 3357.50 (840.78) | 190.63 (7.31) | 281 (7.71) | 528.75 (27.01) |
| **Male** | 15 | 38.79 (1.85) | 3210.67 (421.84) | 193.93 (7.71) | 287 (11.33) | 519.80 (25.03) |
| **Total** | 23 | 38.67 (1.86) | 3269.40 (607.38) | 192.78 (7.148) | 284.91 (10.44) | 522.91 (25.49) |

**Supplementary Table 3**

*Demographical information of the sample for the stability for each one of the pairwise spearman correlations. It displays the mean (SD) of the gestation period, weight at birth and age in each session for female and male separated.*

| | | | Gestation | | Session Age (days) | | |
|---|---|---|---|---|---|---|---|
| | | *n* | *Weeks* | *Weight (grams)* | *1st Session* | *2nd Session* | *3rd Session* |
| 6mo – 9mo | Female | 34 | 39.51 (1.46) | 3437. 65 (562.193) | 193.53 (8.37) | 283.39 (8.36) | - |
| | Male | 38 | 39.59 (1.42) | 3343.03 (415.92) | 194.93 (8.36) | 286.39 (8.53) | - |
| 6mo – 16mo | Female | 18 | 39.30 (1.49) | 3323.89 (595.98) | 192.84 (8.98) | - | 519.58 (25.34) |
| | Male | 43 | 39.05 (1.23) | 3230.54 (386.94) | 194.82 (9.33) | - | 517.86 (23.11) |
| 9mo – 16mo | Female | 14 | 39.15 (1.44) | 3390.71 (429.98) | - | 282.43 (6.64) | 526.71 (24.25) |
| | Male | 14 | 39.12 (1.48) | 320.87 (429.98) | - | 285.95 (11.34) | 521.26 (22.74) |

**Supplementary Table 4**

*The goodness of the fit of the model is (R2 index) provided by the FOOOF toolbox for each Session at parietal and frontal clusters. R2 >.95 for It is evaluated with the R-Squared index. All the infants included in the analysis. had, at least, a 0.95 value of fit.*

| | | Parietal | | | Frontal | | |
|---|---|---|---|---|---|---|---|
| | *n* | *Mean (SD)* | *Min.* | *Max.* | *Mean (SD)* | *Min.* | *Max.* |
| **6mo** | 91 | 0.971 (0.008) | 0.950 | 0.989 | 0.968 (0.008) | 0.952 | 0.987 |
| **9mo** | 76 | 0.978 (0.009) | 0.956 | 0.994 | 0.971 (0.009) | 0.952 | 0.989 |
| **16mo** | 44 | 0.982 (0.007) | 0.960 | 0.994 | 0.977 (0.007) | 0.963 | 0.988 |
| **Total** | 91 | 0.976 (0.009) | 0.950 | 0.994 | 0.971 (0.009) | 0.952 | 0.989 |

Note. The information in the table corresponds to the infants who came to the session and did the resting state protocol in the EEG.

**Supplementary Table 5**

*Descriptive statistics of the aperiodic parameters. It shows the mean (standard deviation) in each age and cluster divided by sex.*

| | | | Exponent | | Offset | |
|---|---|---|---|---|---|---|
| | | *n* | *Parietal* | *Frontal* | *Parietal* | *Frontal* |
| **6mo** | *Female* | 40 | 1.26 (0.36) | 0.89 (0.44) | 1.67 (0.26) | 1.46 (0.36) |
| | *Male* | 51 | 1.36 (0.25) | 0.92 (0.38) | 1.71 (0.18) | 1.43 (0.32) |
| **9mo** | *Female* | 35 | 1.52 (0.19) | 1.23 (0.33) | 1.79 (0.17) | 1.63 (0.25) |
| | *Male* | 42 | 1.59 (0.19) | 1.33 (0.17) | 1.78 (0.17) | 1.63 (0.26) |
| **16mo** | *Female* | 17 | 1.71 (0.15) | 1.55 (0.20) | 1.85 (0.12) | 1.82 (0.14) |
| | *Male* | 27 | 1.66 (0.13) | 1.47 (0.23) | 1.83 (0.09) | 1.72 (0.19) |

Note. The table represents only the infants included in the linear mixed models.

**Supplementary Table 6**

*Mean frequency (SD) in Hz of the alpha peak and descriptive parameters of the infants who were included in the repeated measures analysis.*

| | | Mean Age (days) | Mean Frequency (SD) |
|---|---|---|---|
| **6mo** | *Female* | 190.63 (7.31) | 6.55 (0.61) |
| | *Male* | 193.93 (7.03) | 6.63 (0.68) |
| **9mo** | *Female* | 281 (7.71) | 6.59 (0.42) |
| | *Male* | 287 (11.33) | 6.75 (0.46) |
| **16mo** | *Female* | 528.75 (27) | 7.72 (0.60) |
| | *Male* | 519.8 (25.03) | 7.54 (0.59) |

Note. The total sample consisted of 23 infants (female: 15)

**Supplementary Table 7**

*Descriptive statistics of the periodic power (microvolts) and relative power in each frequency band and cluster. It shows the mean (standard deviation) in each age and cluster divided by sex.*

| | | | | **Theta** | | **Alpha** | | **Beta** | | **Gamma** | |
|---|---|---|---|---|---|---|---|---|---|---|---|
| | | | *n* | *Frontal* | *Parietal* | *Frontal* | *Parietal* | *Frontal* | *Parietal* | *Frontal* | *Parietal* |
| **Periodic Power** | *6mo* | Female | 40 | 2.99 (2.35) | 3.33 (2.29) | 0.49 (0.79) | 0.85 (0.97) | -0.048 (0.039) | -0.054 (0.041) | 0.074 (0.052) | -1.776 (0.245) |
| | | Male | 51 | 2.62 (1.83) | 2.94 (1.94) | 0.76 (1.12) | 1.64 (2.19) | -0.038 (0.033) | -0.038 (0.045) | 0.076 (0.055) | -1.783 (0.232) |
| | *9mo* | Female | 35 | 4.09 (3.80) | 4.65 (3.73) | 0.80 (0.77) | 1.63 (1.13) | -0.058 (0.042) | -0.032 (0.035) | 0.071 (0.080) | -1.892 (0.232) |
| | | Male | 42 | 4.25 (4.64) | 4.40 (4.70) | 1.16 (1.08) | 2.22 (1.60) | -0.060 (0.045) | -0.024 (0.049) | 0.065 (0.040) | -1.830 (0.277) |
| | *16mo* | Female | 17 | 2.67 (1.66) | 2.75 (1.21) | 1.13 (0.90) | 2.03 (1.43) | -0.023 (0.034) | 0.016 (0.062) | 0.029 (0.015) | -1.851 (0.179) |
| | | Male | 27 | 3.30 (3.11) | 3.21 (2.45) | 1.31 (0.89) | 2.57 (1.46) | -0.035 (0.044) | 0.026 (0.054) | 0.037 (0.033) | -1.924 (0.200) |
| **Relative Power** | *6mo* | Female | 40 | 0.74 (0.09) | 0.73 (0.09) | 0.18 (0.08) | 0.21 (0.08) | 0.040 (0.023) | 0.036 (0.016) | 0.035 (0.027) | 0.020 (0.012) |
| | | Male | 51 | 0.70 (0.10) | 0.67 (0.09) | 0.22 (0.09) | 0.27 (0.10) | 0.045 (0.020) | 0.038 (0.011) | 0.035 (0.024) | 0.019 (0.009) |
| | *9mo* | Female | 35 | 0.73 (0.08) | 0.69 (0.10) | 0.21 (0.08) | 0.26 (0.09) | 0.038 (0.015) | 0.034 (0.013) | 0.025 (0.017) | 0.015 (0.009) |
| | | Male | 42 | 0.70 (0.10) | 0.65 (0.09) | 0.24 (0.09) | 0.29 (0.08) | 0.041 (0.017) | 0.040 (0.015) | 0.025 (0.018) | 0.018 (0.010) |
| | *16mo* | Female | 17 | 0.69 (0.09) | 0.63 (0.09) | 0.26 (0.09) | 0.30 (0.09) | 0.040 (0.009) | 0.046 (0.010) | 0.015 (0.006) | 0.015 (0.006) |
| | | Male | 27 | 0.67 (0.09) | 0.60 (0.10) | 0.27 (0.09) | 0.34 (0.10) | 0.043 (0.014) | 0.043 (0.012) | 0.018 (0.010) | 0.013 (0.006) |

Note. The table represents only the infants included in the linear mixed models. As the values of the power are especially low in beta and gamma frequency bands, we used three decimals after the dot.

**Supplementary Figure 1**

*The layout of the high-density net employed in the study.*

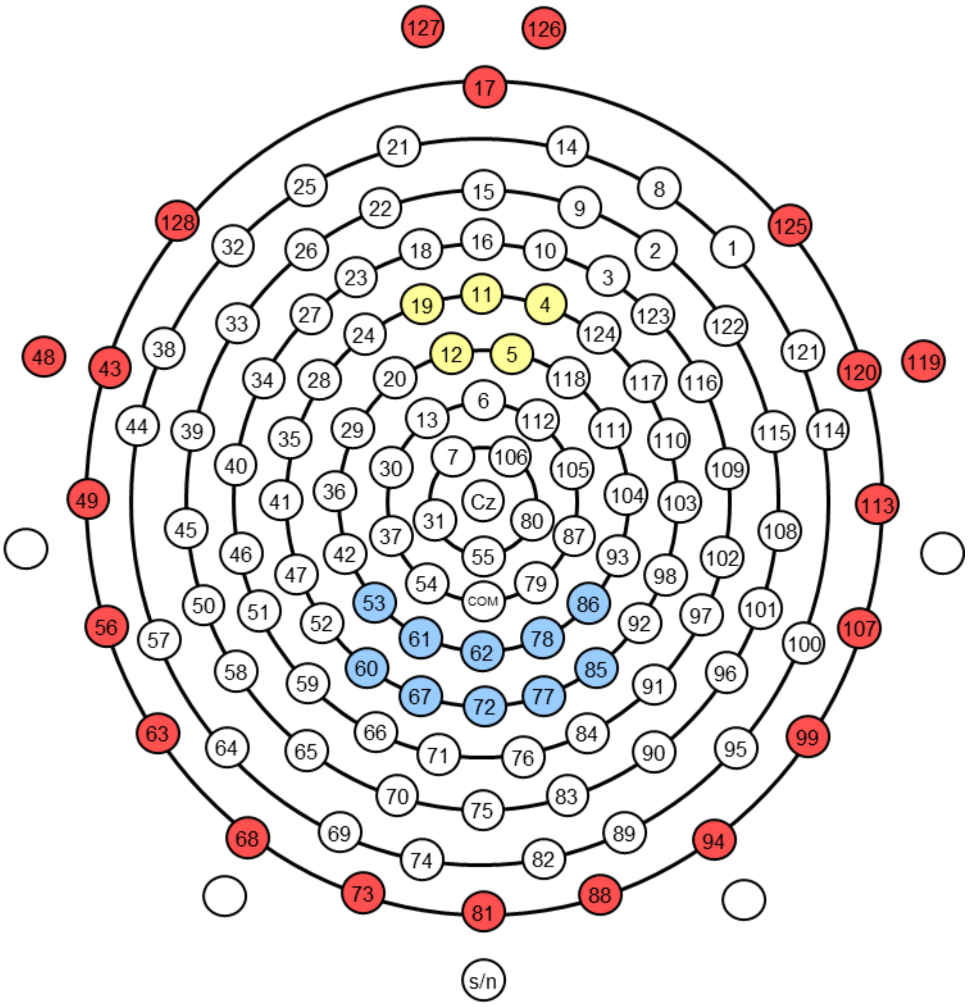


Note. The red colour represents the removed electrodes in the first pre-processing step. The blue and yellow colours represent the electrodes combined in the parietal and frontal clusters, respectively.
