## Supplementary_Results for "Early development of electrophysiological activity: contribution of periodic and aperiodic components of the EEG signal"

As we found the opposite results in the aperiodic parameters in comparison to previous literature, we decided to compute the aperiodic parameters (offset and exponent) changing the frequency range (1 – 10 Hz) as in previous studies with infants. Then, we performed linear mixed models (LMM) analysis for each parameter including Session-Age (6, 9 and 16 months) and Cluster (Parietal and Frontal) as fixed effects. Additionally, we included the participant as a random effect to account for individual variability. See Supplementary Tables 8 for information about the fit of the aperiodic component and Supplementary Figure 2 for the aperiodic curves computed.

**Supplementary Table 8**

*The goodness of the fit of the model is provided by the FOOOF toolbox in the 1 to 10 Hz. It is evaluated with the R-Squared index. All the infants included in the analysis had, at least, a 0.95 value of fit.*

| | | **Parietal** | | | **Frontal** | | |
|---|---|---|---|---|---|---|---|
| | *n* | *Mean (SD)* | *Min.* | *Max.* | *Mean (SD)* | *Min.* | *Max.* |
| **6mo** | 91 | 0.994 (0.005) | 0.972 | 0.989 | 0.993 (0.005) | 0.968 | 0.989 |
| **9mo** | 76 | 0.991 (0.006) | 0.974 | 0.999 | 0.988 (0.008) | 0.967 | 0.998 |
| **16mo** | 44 | 0.989 (0.007) | 0.965 | 0.998 | 0.988 (0.007) | 0.965 | 0.998 |
| **Total** | 91 | 0.992 (0.006) | 0.965 | 0.998 | 0.991 (0.007) | 0.965 | 0.998 |

Note. The information in the table corresponds to the infants who came to the session and did the resting state protocol in the EEG.

**Supplementary Figure 2**

*Aperiodic curves for the 1 to 10 Hz range (A) and the 1 to 45 Hz range (B)*

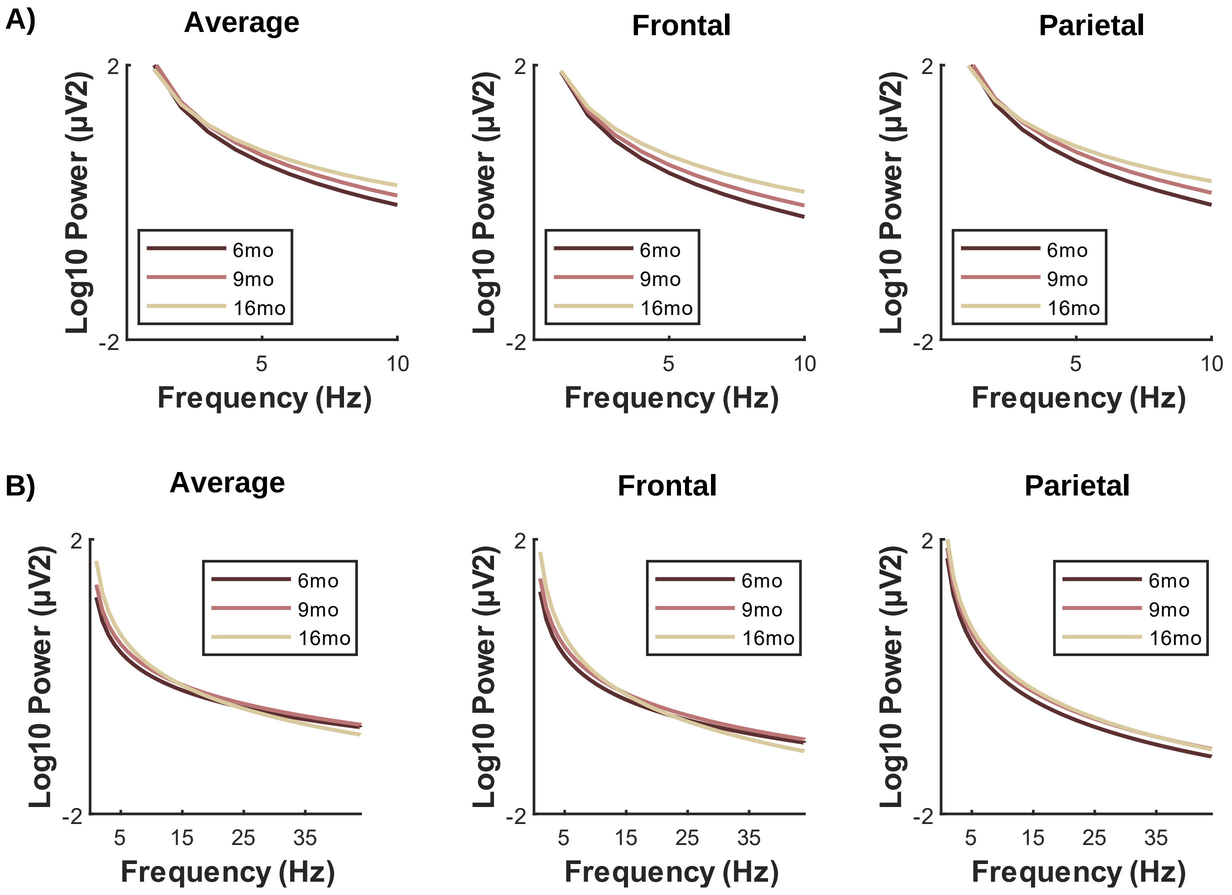


Note. It displays the curves for the average of all electrodes and the parietal and frontal clusters divided by session age.

The offset was larger in the parietal cluster ($F$(1,213.11) = 306, $p$ < .001) and changed across sessions ($F$(2,316.85) = 4.76, $p$ = .009; Supplementary Figure 3A) but the interaction between Cluster and Session Age was not significant ($F$ = 1). The EMs of the session age revealed higher values of the offset at 9mo in comparison to 6mo ($p$ = .018) and 16mo ($p$ = .034) sessions but no differences between 6mo and 16mo ($p$ = 1).

The exponent had higher values in the parietal cluster ($F(1,221.12) = 15.43$, $p < .001$) and varied across sessions ($F(2,317.32) = 87.86$, $p < .001$; Supplementary Figure 3B) but both factor did not interact ($F < 2$). The exponent descended from the first to the second session ($p < .001$) and from the second to the third one ($p < .001$).

**Supplementary Figure 3**

*Aperiodic components develop in the 1 to 10 Hz frequency range.*

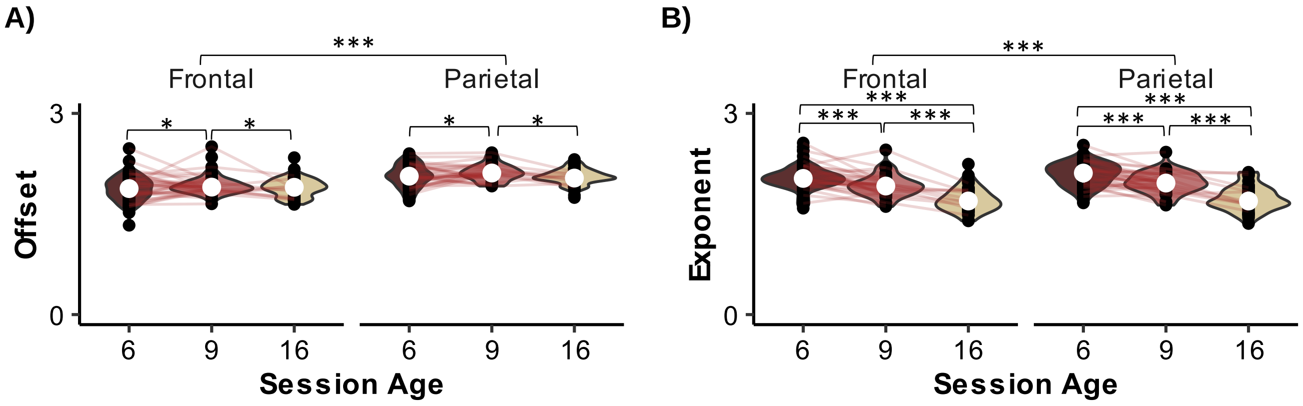


Note. It displays a violin plot for each aperiodic component (offset and exponent) across sessions divided by cluster. Black dots represent the individual values of each infant, whereas the white dot and white line correspond to the median and the first and third quartiles, respectively. The red lines that unite the black dots are the trajectory of each infant.*$p < .05$, ** $p < .01$, *** $p < .001$.

To determine whether the offset and exponent values differed between the 1 to 10 Hz and 1 to 45 Hz range, we decided to compute in a linear mixed model including the age and frequency range and fixed effects and their interaction for each cluster separately. As we have described the trajectories of the aperiodic components in the main text and supplementary material for each frequency range, in the following paragraphs we focused on the frequency range main effect and the interaction.

In the frontal cluster, the offset was larger in the 1 to 10 Hz range ($F(1, 185.07) = 220.76$, $p < .001$) which interacted with the age ($F(2, 202.753) = 21.49$, $p < .001$). The EMs revealed that the narrowest range always displayed a higher offset (*p*s < .003). Similarly, at the parietal cluster the 1 to 45 Hz had lower offset values ($F(1, 200.35) = 208.347$, $p < .001$) and the interaction was also significant ($F(2, 205.36) = 47.44$, $p < .001$). The planned comparisons showed augmented values of the offset in the 1 to 10 Hz in the first two sessions (*p*s < .001) but no differences in the third session ($p = .393$). See Supplementary Figure 4A.

In relation to the exponent, at the frontal cluster, the 1 to 10 Hz had taller values ($F(1, 160.30) = 222$, $p < .001$) and it interaction with the session age ($F(2, 196.924) = 52.82$, $p < .001$). The exponent values were higher only in the two first sessions (*p*s < .001) for the narrowest frequency band but the values did not vary in the third session ($p = .662$). At the parietal cluster, again the 1 to 10 Hz had augmented values ($F(1,153.85) = 41.26$, $p < .001$) which interacted with the session age ($F(1, 195.07) = 71.94$, $p < .001$). In this case, the parietal cluster had always larger exponent in all the sessions (*p*s < .001). See Supplementary Figure 4B.

**Supplementary Figure 4**

*Offset and Exponent variations between frequency ranges (1 to 10 Hz and 1 to 45 Hz) within each session and cluster.*

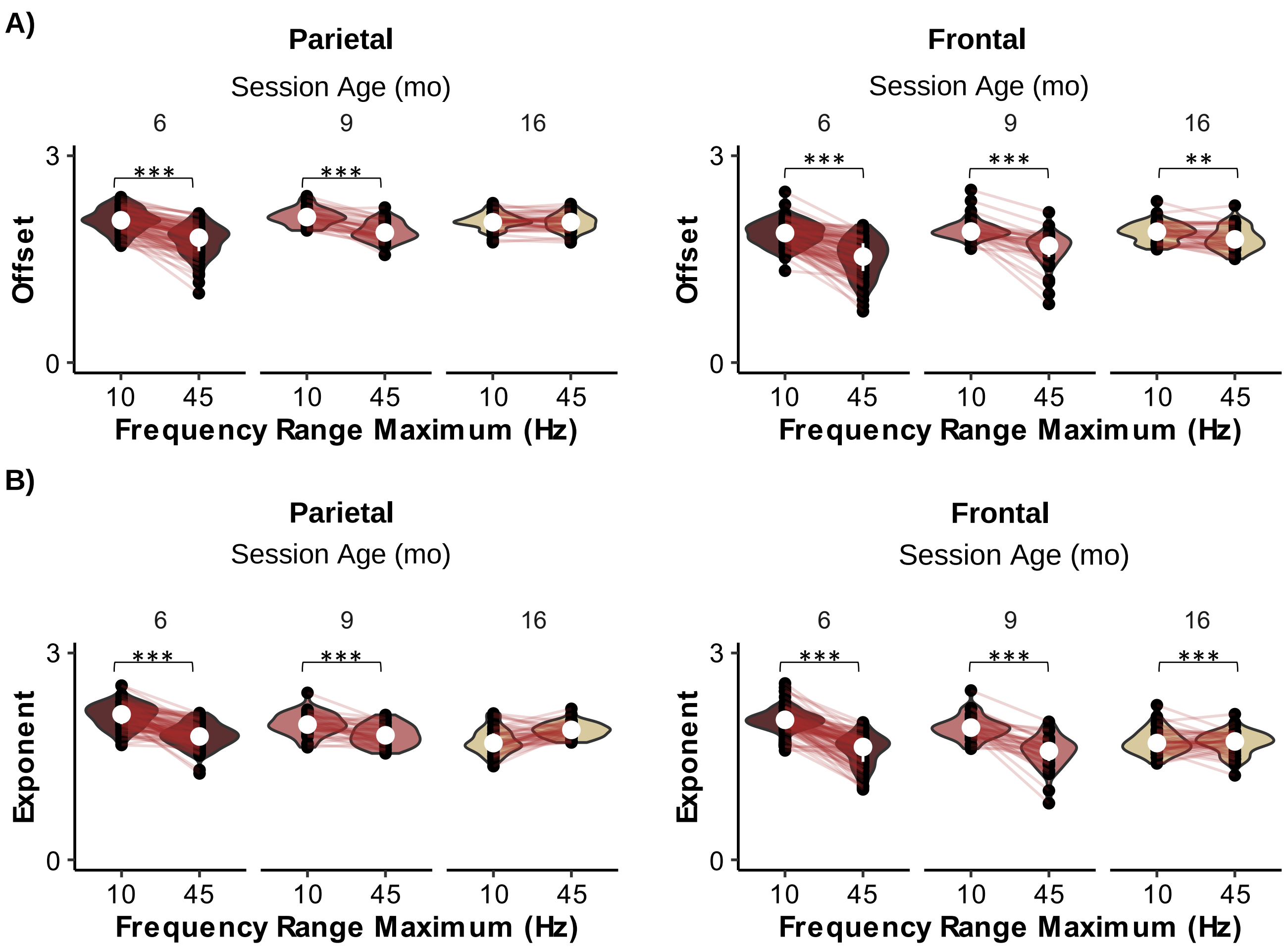


Note. It displays a violin plot for each aperiodic component (offset and exponent) comparing the two ranges employed (1 to 10 Hz and 1 to 45 Hz) divided by session and cluster. Black dots represent the individual values of each infant, whereas the white dot and white line correspond to the median and the first and third quartiles, respectively. The red lines that unite the black dots are the trajectory of each infant. ** $p < .01$, *** $p < .001$.
